# NPY+ Interneurons in Basolateral Amygdala are Activated by Aversive Stimuli

**DOI:** 10.64898/2026.04.30.722047

**Authors:** Patric J. Perez, Aundrea F. Bartley, J. Andrew Hardaway, Lynn E. Dobrunz

## Abstract

Traumatic events increase the risk for anxiety disorders, yet knowledge of how trauma modulates neuronal activity to induce anxiety is incomplete. The amygdala, which processes stressful sensory information, is enriched with interneurons that release the anxiolytic neurotransmitter neuropeptide Y (NPY). Amygdala NPY levels are reduced one week after an aversive event, suggesting chronic alteration of NPY+ interneurons; however, studies of *in vivo* amygdalar NPY+ cell activity during stressors are lacking. Here, we use a genetically encoded calcium sensor together with fiber photometry to investigate *in vivo* activation of NPY+ cells in basolateral amygdala (BLA) to aversive stimuli in mice. NPY+ cell activation was evaluated in response to two aversive stimuli, air puffs to the face (mild) and footshocks (strong). Air puffs caused a transient elevation of calcium in BLA NPY+ cells, indicating robust neuronal activation, in both male and female mice with no sex-dependent differences. Interestingly, there was habituation of the calcium signal in NPY+ cells to later air puff iterations. Strong footshocks also caused calcium elevation in both male and female mice with no sex-dependent differences. Excitingly, footshock induces a larger calcium response compared to air-puff. In contrast to air puff, the calcium signal to footshock was prolonged in later iterations. BLA NPY+ cell calcium signals were consistent in response to the same footshock protocol delivered 1 week later, indicating that activation of NPY+ cells by footshock is stable across this timeframe. Taken together, these results reveal a potential role for NPY+ interneurons in basolateral amygdala during aversive events.

## 1. INTRODUCTION

The amygdala is a critical integrator of fear learning (Johansen et al., 2011; Maren & Quirk, 2004; Pape & Pare, 2010) and modulates responses to stress (Davis & Whalen, 2001; Janak & Tye, 2015; Jie et al., 2018) and aversive events (Sengupta et al., 2018; X. Zhang et al., 2021). Specifically, basolateral amygdala (BLA) can discriminate between aversive stimuli (Corder et al., 2019; Grosso et al., 2018; Liu et al., 2021) and mediate appropriate behavioral responses (Gore et al., 2015; Sparta et al., 2014). Stressful events induce long-term changes in amygdala, including neural circuit hyperactivity (Koenigs & Grafman, 2009; Rauch et al., 2000; Shin et al., 2006), potentially due to disruptions in interneuron activity (Jie et al., 2018; Martijena et al., 2002; Perumal & Sah, 2021). Animal models of stress consistently demonstrate chronic alterations in GABAergic inhibition, evidenced by decreases in inhibitory synaptic transmission (Isoardi et al., 2007) and GAD65/7 expression (Tzanoulinou et al., 2014). However, information on the real-time responses of GABAergic interneurons to varying degrees of aversive stimuli in amygdala is incomplete.

A subset of GABAergic interneurons in amygdala expresses neuropeptide Y (NPY) (Hájos, 2021; Mańko et al., 2012; Ozsvár et al., 2024; Vereczki et al., 2021), a key anxiolytic neuropeptide in the brain (Broqua et al., 1995; Christiansen et al., 2014; Heilig et al., 1989). In BLA, NPY+ interneurons are 15% of GABAergic cells (Hájos, 2021), suggesting a potential role in aversive stimuli discrimination. NPY receptor agonists injected into amygdala reduce anxiety-like behavior (Gutman et al., 2008; Heilig et al., 1993; Kornhuber & Zoicas, 2021; Tammy J Sajdyk et al., 2006), and NPY injection into BLA induces resilience (Michaelson et al., 2020; Silveira Villarroel et al., 2018). Blocking or knocking down NPY receptors in amygdala produces anxiogenic effects (Heilig, 1995; T J Sajdyk et al., 1999), highlighting the importance of endogenous NPY release in amygdala for regulating anxiety. Stress paradigms can induce lasting changes in NPY mRNA levels (Reichmann & Holzer, 2016; Thorsell et al., 1998), NPY expression (Cohen et al. 2012; Cohen et al. 2015; Cortes et al. 2021), or NPY release (Li et al. 2017), suggesting that aversive stimuli can directly affect NPY+ cells. Consistent with this, juvenile restraint stress increased expression of the activity marker c-Fos in hippocampal NPY+ cells (Regev-Tsur et al., 2020). However, elevated plus maze failed to increase c-Fos expression in the amygdala (Butler et al., 2012), indicating that not all aversive stimuli activate NPY+ interneurons. Our understanding of the effects of different types of stressors on BLA NPY+ cells is incomplete, and the dynamics of *in vivo* BLA NPY+ cell activation during aversive stimulation remain uninvestigated.

Here we measured *in vivo* NPY+ cell activation using fiber photometry and Cre-dependent GCaMP8f in NPY-Cre mice. Our results demonstrate for the first time that BLA NPY+ interneurons show robust calcium transients during both air puff and footshock, with only subtle differences, indicating that they are acutely activated by both mildly and strongly aversive stimuli in freely behaving mice.

## 2. MATERIALS AND METHODS

### 2.1 Animals

All experimental procedures were carried out in compliance with the Institutional Animal Care and Use Committee of the University of Alabama at Birmingham. Experiments were conducted in accordance with the National Institutes of Health Guide for the Care and Use of Laboratory Animals. Adult male and female Tg(Npy-cre)RH26Gsat (Gong et al., 2007) (RRID: MMRRC_037423-UCD; part of the GENSAT project) Cre+/− mice, aged 13-23 weeks, were used for all *in vivo* fiber photometry experiments. All mice were group-housed, 3 to 7 in a cage, while food and water were provided *ad libitum.* Experiments were performed from 8 AM to 6 PM. Mouse colonies were maintained on a 12-hour light/dark cycle.

### 2.2 Viral Vectors

Adeno-associated viruses (AAV) encoding GCaMP8f (pGP-AAV-syn-FLEX-jGCaMP8f-WPRE #162379) were purchased from Addgene. Viral aliquots were diluted with sterile phosphate buffered saline (PBS) to reach the titer used for injection (GCaMP8f: 6 x 10^12^ GC/m).

### 2.3 Stereotaxic Surgery

Stereotaxic transcranial surgery was performed for viral injections and fiber optic canula insertions.

#### 2.3.1 Viral injections

All instruments were sterilized before being used on animals. 8-11 week old mice were induced at 5% using an isoflurane vaporizer, which was slowly tapered down to 1% until the cessation of the surgery. Additionally, after induction, a drug cocktail consisting of buprenorphine and meloxicam (0.1 mg/kg and 5 mg/kg, respectively) was subdermally injected. A Vaseline-coated rectal probe (Harvard Instruments, USA) was inserted to ensure a body temperature of 36 °C was maintained. The head is fixed using the ear bars and incisor bar on the stereotaxic apparatus (Kopf Instruments, USA). Hair was removed from the scalp with hemostats and aseptically cleaned with 3 washes of iodine and 70% ethanol, along with an application of emulsified 5% lidocaine and triple antibiotic ointment before the dermal incision. The brain was leveled to within 100 µm in both the anterior/posterior & medial/lateral axes using bregma and lambda. Bilateral craniotomies were drilled over the basolateral amygdalae (BLA; A/P: –1.45, M/L: ±3.15, D/V: –4.75 from Bregma) for viral injection. 300 nL of virus per hemisphere were injected at a rate of 100 nL/minute and the Hamilton syringe remained in the brain for 5 minutes post-injection.

#### 2.3.2 Insertion of fiber optic cannulas

Before cannula insertion, three small holes were drilled to place head anchor screws. Fiber photometry experiments used fiber optic cannulas (200 µm diameter; Xi’an HealthiGlobal Tech, China) inserted bilaterally into the amygdala. Cannulas were lowered into the amygdalae and secured with dental cement (Stoelting, USA). Postoperative care consisted of a 3% (w/v) subdermal saline or Ringer’s injection immediately after surgery, along with 0.1 mg/kg buprenorphine injections every 12 hours for two days and 5 mg/kg meloxicam injection every 24 hours for two days. All mice were allowed to recover for at least 3 weeks before any handling. Fiber photometry recordings occurred 4 to 8 weeks after surgery.

#### 2.3.3 Histology

After completion of experiments, histology was performed to confirm fiber placement and viral expression. Mice were heavily anesthetized with 5% isoflurane. A priming solution containing 1X PBS with 10 U of heparin per mL was perfused into the animals to prevent blood clotting and aid in paraformaldehyde infiltration into the brain microvasculature. Then, perfusion with 4% paraformaldehyde in 1X PBS, pH = 7.4, was used for fixation. Brains were excised and post-fixed for 24 hours. Afterward, the tissue was placed in 30% sucrose for cryopreservation. Brains were sectioned in a cryostat (Leica, Germany) at 40 µm and placed into a 24-well plate in typewriter fashion to maintain slice order. Fiber placement was verified by directly mounting the tissue onto slides. All slides were mounted with Glass Antifade Mountant (Invitrogen, USA), imaged, and later sealed with clear nail polish. Imaging was done on a Keyence BZ-X810 (Japan). Briefly, a 20X objective (NA = 0.75) was used to tile stitch together a region of interest, and the focal plane was chosen based on the emission spectra of the fluorophore. Hemispheres were included if the majority of GCaMP expression was observed in the BLA; some hemispheres also had minor GCaMP expression in the central nucleus of the amygdala. Hemispheres were excluded if: 1) no detectable expression of GCaMP was observed in BLA 2) the fiber track/tip was not found, or 3) the fiber tip was more than 1 mm away from the cluster of cells expressing GCaMP. Three hemispheres were excluded based on histology criteria.

### 2.4 In Vivo Fiber Photometry

#### 2.4.1 Handling of mice

All mice were handled for at least one 5-10 minute session each day over three days before the start of fiber photometry experiments. Handling included transportation from the vivarium to the laboratory, acclimation to the behavior room for 30 minutes, scruffing, tethering to the patch cables (10 min), and acclimation to the behavior box (10 min). The clear plastic box (dimensions 20.32 x 30.48 x 23.49 cm) was cleaned with deionized water between handling and experimental sessions.

#### 2.4.2 Fiber photometry

The FP3002 fiber photometry system (Neurophotometrics, USA) was used for all experiments. For this setup, both the 470 nm and 415 nm LEDs’ excitation wavelengths were interleaved and transmitted through a 200 μm core, 0.37 NA patchcord with a tip excitation power of 50 μW. To accurately align stimuli to neuronal activity, TTL-generated timestamps were collected by interfacing an Arduino with Bonsai software (Lopes et al., 2015). Additionally, the Thor ADAL3 (Thor Labs, USA) connector was used to attain a robust connection between the fiber optic cannula and the patch cable. The fiber optic cannulas were custom manufactured to 5.5mm and had > 80% light transmission (#FOC-C-B-200-1.25-0.37-5.5, Xi’an HealthiGlobal Tech, China). For each experiment, a recording period of 30 minutes was collected to photobleach the patch cable and serve as the acclimation period.

#### 2.4.3 Fiber photometry data analysis

All photometry data were collected as comma separated value (csv) files. Data for the 470 nm (signal) and 415 nm (reference) wavelengths were collected at 20 frames per second per wavelength. Timestamps were recorded for each event (air-puff or footshock) iteration. Data was processed using an established fiber photometry pipeline, pMAT (Bruno et al., 2021) which creates a scaled reference curve, calculates ΔF/F, and converts the results to Z-score traces. Data were analyzed using a bin constant of 4, resulting in 0.2 s / point. Traces for each event were time locked using a 90 s window starting 30 s before the stimulus. Z-score traces for each trial were vertically translated in Origin to remove baseline offset prior to the stimulus. For each hemisphere, the traces for each event were averaged together, unless otherwise noted. The peak and area under the curve (AUC) were calculated from the average Z-score traces per hemisphere. Hemispheres were classified as non-responsive if the Z-score peak of the average trace was below 1. Two hemispheres from one animal were non-responsive to both air puff and footshock stimuli. Additionally, these hemispheres had low expression of GCaMP, indicating that the GCaMP signal detected by the fiber was likely insufficient.

#### 2.4.4 Air Puff

The air puff (compressed gas duster) was presented through a tube placed < 5 cm from the face of the animal. Mice were exposed to 5 interspaced brief (< 2 seconds) air puffs; each exposure was at least 5 minutes apart. Each time stamp was added using a push-button switch. Two hemispheres from one animal were classified as non-responsive and removed from the data set.

#### 2.4.5 Footshock

A footshock box (Med Associates, USA) was used to deliver 10 regularly interspaced footshocks (1 milliampere, 1 second) which were applied every 5 minutes in a shock chamber (VFC-008) of dimensions 29 x 24 x 21 cm via a shock-grid floors (VFC-005A). The light intensity in the footshock box was 230 lux. Three hemispheres (2 from one animal and one from another) were classified as non-responsive and removed from the data set.

### 2.5 Behavior

The open field (OF) behavioral task was used to measure locomotor activity and thigmotaxis 7 days after exposure to the footshock protocol using NPY-Cre^-/-^ mice. The OF apparatus is a square (44 cm^2^) box made from plexiglass. All OF experiments were conducted with light levels around 30 lux and lasted 5 minutes. EthoVision (Noldus, USA) software was used to track the total ambulatory time, distance traveled, and velocity in each zone (center and periphery) of the OF apparatus. The percentage of distance traveled in the center region of the OF was calculated.

### 2.6 Statistics

Data are reported as mean ± standard error (SEM). OriginPro 2024/5 (USA) was used to detect differences with one– or two-tailed paired t-tests or two-tailed Student’s t-tests. For behavior, n numbers were based on individual animals. For fiber photometry, n numbers were based on individual hemispheres, with number of animals also reported.

## 3. RESULTS

### 3.1 BLA NPY+ Interneurons Respond to Aversive Air Puff Stimuli

Recently, multiple types of amygdalar GABAergic interneurons have been shown to respond to aversive stimuli *in vivo* (Asim et al., 2024) however GABAergic interneurons that release the anxiolytic NPY have not yet been fully evaluated. First, we used an air puff to the face, which is a mild aversive stimulus across species, including humans (Kauvar et al., 2025) that has previously been shown to activate a subpopulation of BLA neurons (Zhang and Li 2018; Zhang et al. 2021). We measured activation of NPY+ interneurons in the BLA using fiber photometry to record acute population-level calcium activity in transgenic NPY-Cre^+/−^ mice injected with cre-dependent GCaMP8f (Figure 1A). Bilateral injections and fiber placement targeted the BLA (Figure 1B), resulting primarily in GCaMP expression and fiber tips in BLA (Figure 1C). Air puff caused a transient increase in GCaMP fluorescence from NPY+ cells (Figure 1D). Both peak Z-scores (Figure 1E) and the area-under-the-curve (AUC, Figure 1F) were significantly higher during air puffs than during the preceding baseline period. Maximal activation of BLA NPY+ neurons occurs quickly as the latency to peak fluorescence ranged from 0.1 to 1.5 seconds (average: 0.52 ± 0.14 s). It appears that the approach/looming by the researcher before the air puff delivery might cause minor activation of BLA NPY+ cells, as evidenced by the small rise in the calcium signal starting approximately 4 seconds before the air puff exposure (t = 0). Peak activation persisted across multiple air puff trials (Figure 1G), and no difference in the peak Z-score was observed between the first and fifth air puff (Figure 1H). Interestingly, the AUC (Figure 1I) and the half-width (Figure 1J) of the calcium signal diminish on the fifth air puff iteration compared to the first air puff, suggesting habituation of the NPY+ cells to repeated exposure. Together, these data confirm that a mild aversive stimulus activates NPY+ neurons in BLA, and the duration of the calcium signal diminishes with repeated trials.

**Figure 1.**
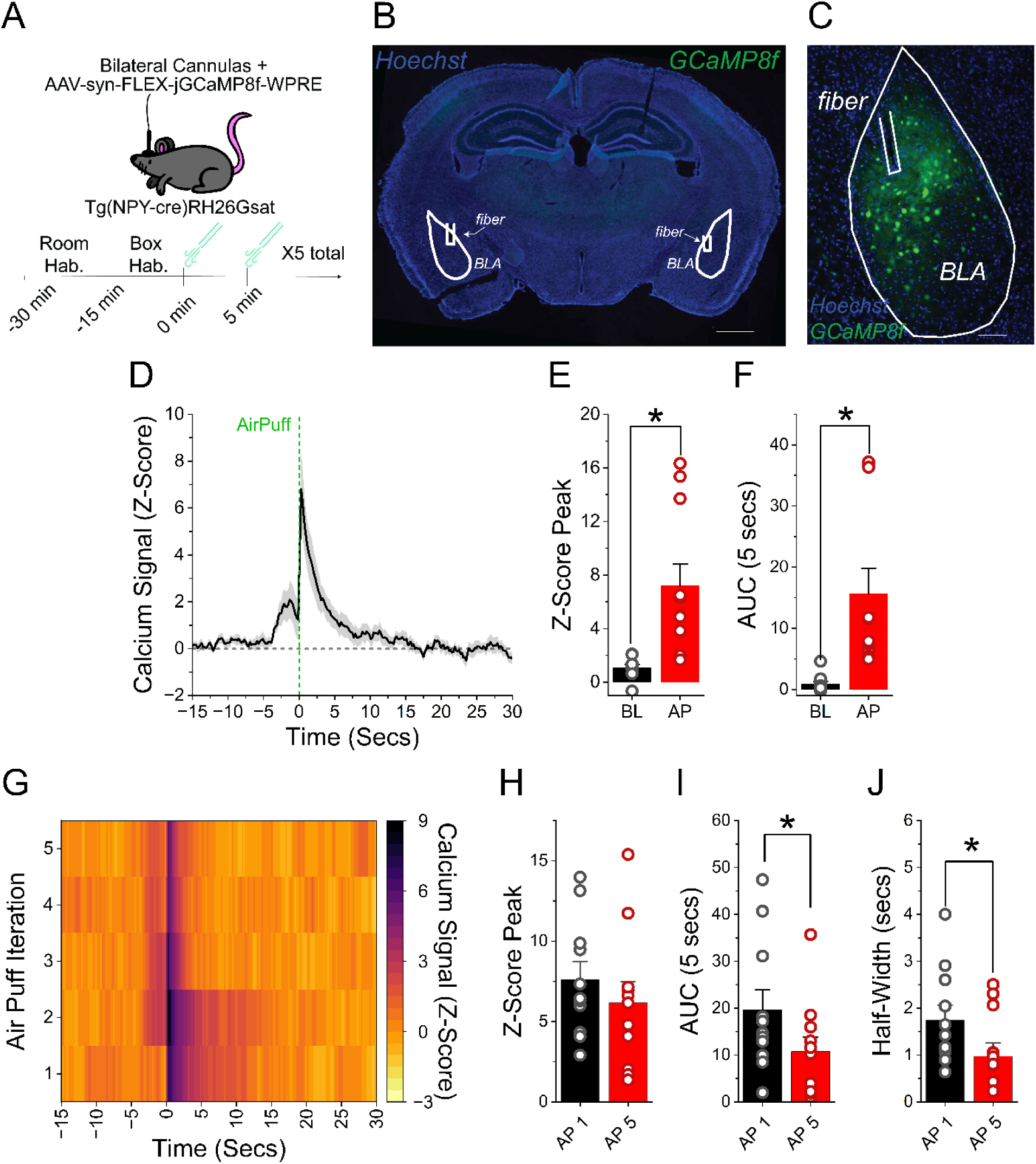
BLA NPY+ Interneurons Respond to Air Puff Stimulus. A. Surgical and experimental design. Bilateral AAV injections of Cre-dependent GCaMP8f followed by fiber optic cannula insertion were performed in male and female NPY-Cre^+/−^ mice to target NPY+ interneurons in the BLA. The aversive stimuli consisted of 5 air puffs (at least 5 minutes apart). B. Representative image showing bilateral fiber placement into the BLA. Scale bar = 1 mm. C. Higher magnification image of transduced NPY cells with fiber placement. Scale bar = 250 μm. D. GCaMP fluorescence shows increased calcium response (n = 11 hemispheres/7 mice), indicating that air-puff activates BLA NPY+ cells. E. Peak Z-Score is significantly larger during the air-puff (AP) compared to baseline (BL; Two-tailed paired *t*-test, p<0.003). F. Area under the curve (AUC, 0 to 5 seconds) is greater during air-puffs compared to baseline (–15 to –10 seconds; Two-tailed paired *t*-test, p<0.006). G. Heat map displaying the Z-Score value of each air-puff iteration averaged across all hemispheres, where t = 0 is air-puff application. H. Peak Z-Score is not significantly lower for the fifth iteration compared to the first iteration of air-puffs (One-tailed paired *t*-test, p=0.90). I. AUC (0 to 5 seconds) is significantly lower between the first and fifth iterations of air-puffs (One-tailed paired *t*-test, p<0.007), suggesting habituation of BLA NPY+ cell activation to repeated air-puffs. J. The calcium signal half-width narrows on the fifth air-puff iteration compared to the first (One-tailed paired *t*-test, p<0.008), suggesting a shorter duration of NPY+ cells activation. For all panels the error bars represent ± SEM. * indicates *p* < 0.05.

### 3.2 BLA NPY+ Interneurons are Activated by Air Puff in both Male and Female Mice

Substantial sex differences in activity of amygdala neurons have previously been reported (Blume et al., 2017). Here sex-specific analyses showed that both female and male mice exhibited significant increases in GCaMP signals from BLA NPY+ cells upon air puff exposure (Figures 2A-B). However, there were no sex differences in Z-Score peak values (Figure 2C), AUC (Figure 2D-E), or half-width of the calcium signal (Figure 2F). Collectively, these data demonstrate that a mild aversive stimulus can activate BLA NPY+ neurons in both sexes.

**Figure 2.**
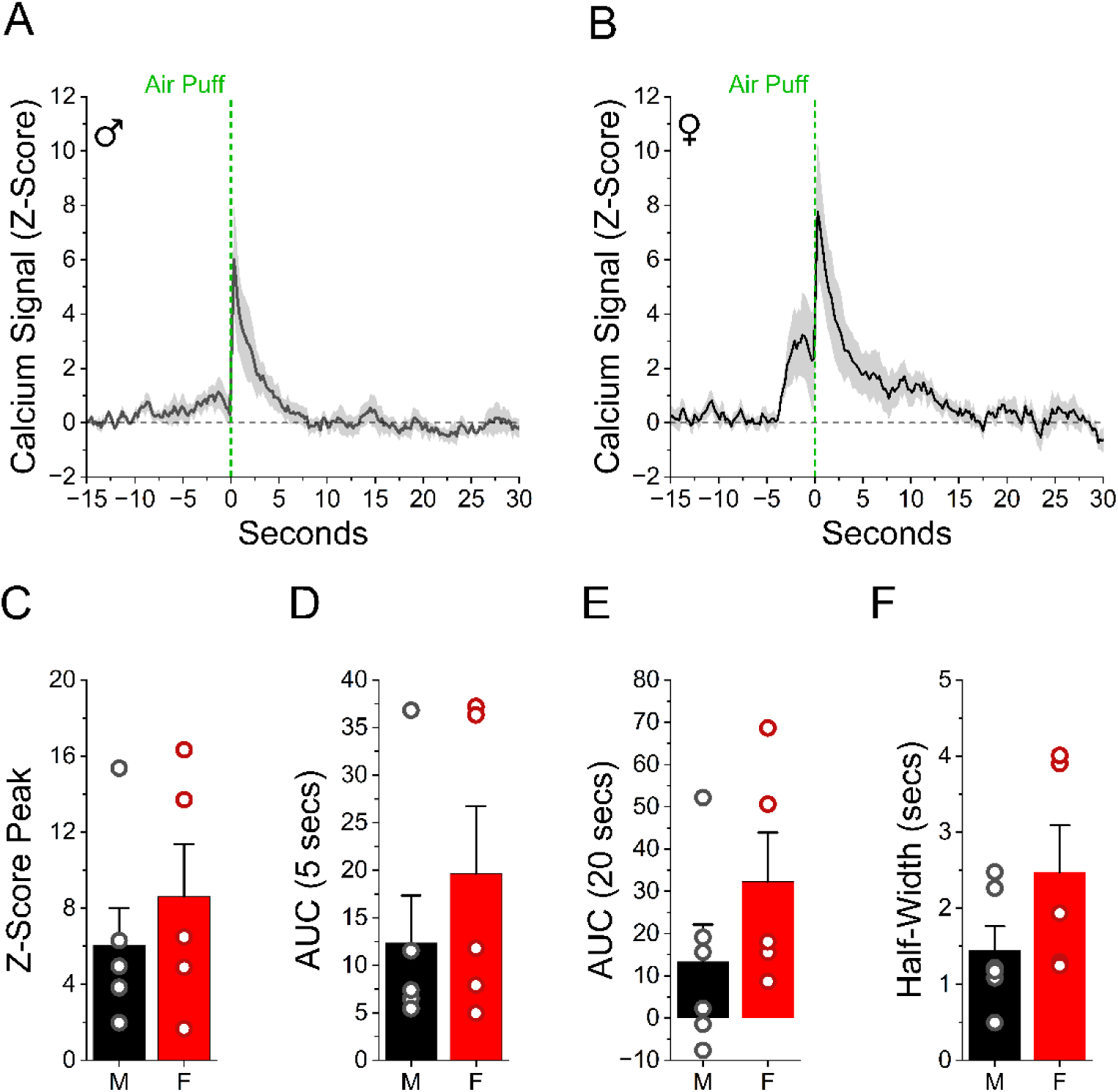
BLA NPY+ Interneurons Respond to Air Puff Stimulus in Both Males and Females. A. Calcium fiber photometry trace shows air-puff activates NPY cells in male mice (n= 6 hemispheres/4 mice). B. Calcium fiber photometry trace shows air-puff activates NPY cells in female mice (n= 5 hemispheres/3 mice). C. Peak Z-score is not different between female and male mice. (Two-tailed Student’s *t*– test, p=0.46). D. AUC (0 to 5 seconds) is not different between female and male mice. (Two-tailed Student’s *t*-test, p=0.41). E. Even a longer AUC (0 to 20 seconds) shows no difference between female and male mice. (Two-tailed Student’s *t*-test, p=0.22). F. The half-width of the calcium signal is not different between male and female mice. (Two-tailed Student’s *t*-test, p=0.15). For all panels the error bars represent ± SEM. * indicates *p* < 0.05.

### 3.3 BLA NPY+ Interneurons Respond to Aversive Footshock Stimuli

In the BLA, some neurons respond to multiple types of aversive stimuli, and others respond only to a specific stimulus (Corder et al., 2019; Liu et al., 2021). It is unknown if different stimuli will activate NPY+ interneurons in BLA. Here, we used footshock (Figure 3A), a strong aversive stimulus well known to activate various cell types in the amygdala (Hochgerner et al., 2023; Sengupta et al., 2018). The footshock protocol (1 sec duration, 1 mA, average of 10 trials, Figure 3A) induced a robust and transient increase in the calcium signal from NPY+ cells (Figure 3B). Both peak Z-scores (Figure 3C) and the AUC (Figure 3D) were significantly higher during footshock as compared to the preceding baseline period. BLA NPY+ neurons reach a maximal peak activation quickly, as the average latency to peak is 0.72 ± 0.08 s. Activation persisted across repeated footshock trials (Figure 3E), and there was no difference in the peak Z-score between the first and last footshock (Figure 3F), indicating that NPY+ cells are reliably activated across all 10 trials. Excitingly, the duration of the calcium signal was prolonged with later iterations of the strong aversive stimuli, as reflected by an increase in the AUC (Figure 3G) and half-width (Figure 3H) on the tenth footshock trial compared to the first. Together, these data confirm that footshock rapidly and repeatedly activates NPY+ neurons in BLA.

**Figure 3.**
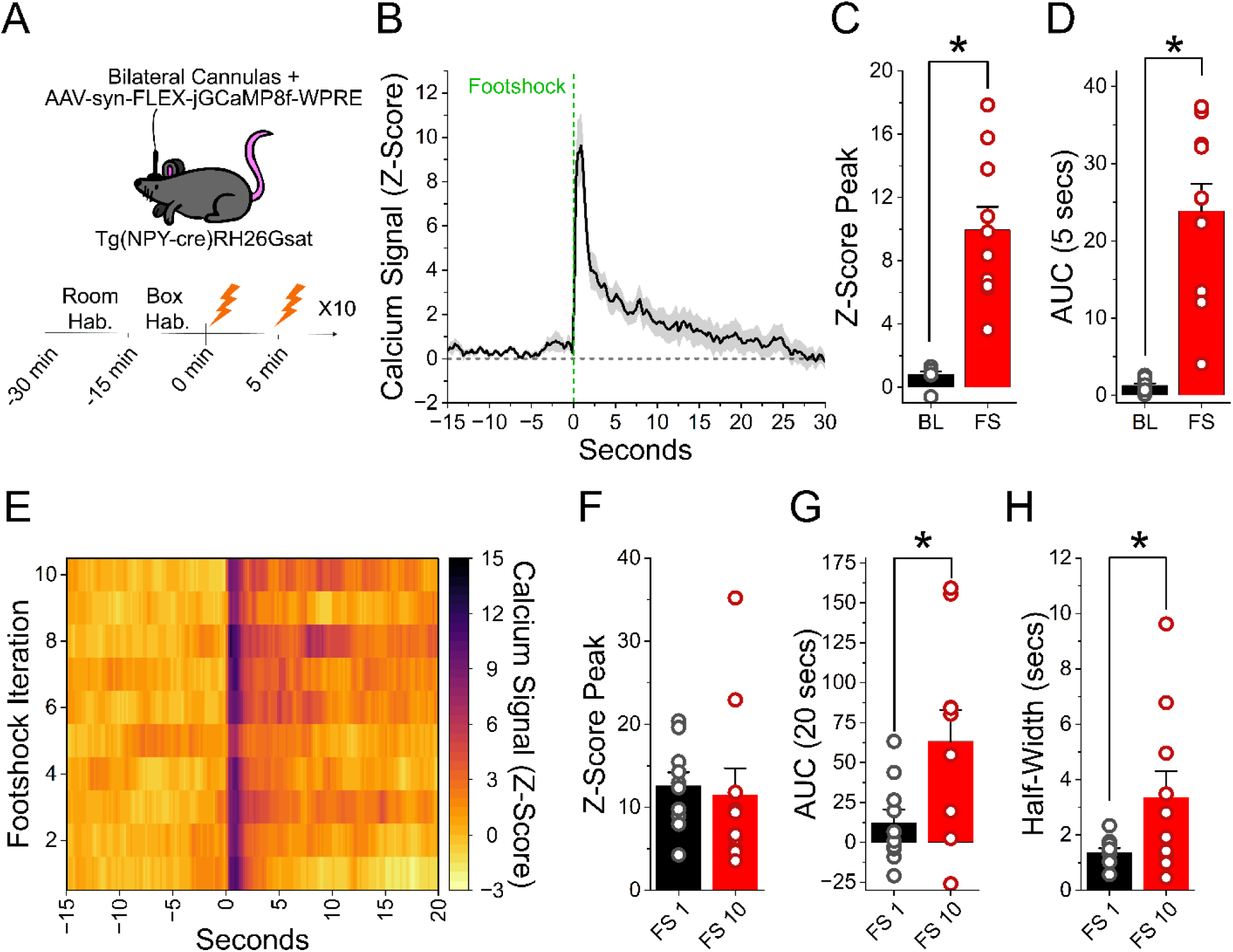
BLA NPY+ Interneurons Respond During Footshock Stimulus. A. Surgical and experimental design. Bilateral AAV injections of Cre-dependent GCaMP8f followed by fiber optic cannula insertion were performed in male and female NPY-Cre^+/−^ mice to target NPY+ interneurons in the amygdala. The aversive stimuli consisted of 10 footshocks (1 sec, 1 mA, 5 minutes apart). B. GCaMP fluorescence shows increased calcium response (n = 10 hemispheres/7 mice), indicating that footshock activates NPY+ cells. C. Peak Z-Score is significantly larger in response to the footshock (FS) compared to baseline (BL; Two-tailed paired *t*-test, p<0.005). D. Area under the curve (AUC, 0 to 5 seconds) is greater for footshock compared to baseline (–10 to –5 seconds; Two-tailed paired *t*-test, p<0.005). E. Heat map displaying the Z-Score value of each footshock iteration averaged across all hemispheres, where t = 0 is footshock application. F. Peak Z-Score is not significantly different between first and tenth iterations (One-tailed paired *t*-test, p=0.66). G. AUC (0 to 20 seconds) is larger for the tenth footshock iteration compared to the first (One-tailed paired *t*-test, p=0.017). H. The calcium signal half-width broadens on the tenth footshock iteration compared to the first (One-tailed paired *t*-test, p=0.024), suggesting prolonged duration of BLA NPY+ cell activation. For all panels the error bars represent ± SEM. * indicates *p* < 0.05.

### 3.4 BLA NPY+ Neurons are Activated by Footshock in both Male and Female Mice

The calcium signal from BLA NPY+ cells was enhanced in both sexes with footshock delivery (Figures 4A-B). However, no difference between males and females were observed for Z-Score peak values (Figure 4C), AUC (Figure 4D-E), or half-width of the calcium signal (Figure 4F). Collectively, these data demonstrate that strong aversive stimuli produce robust activation of BLA NPY+ neurons in both male and female mice.

**Figure 4.**
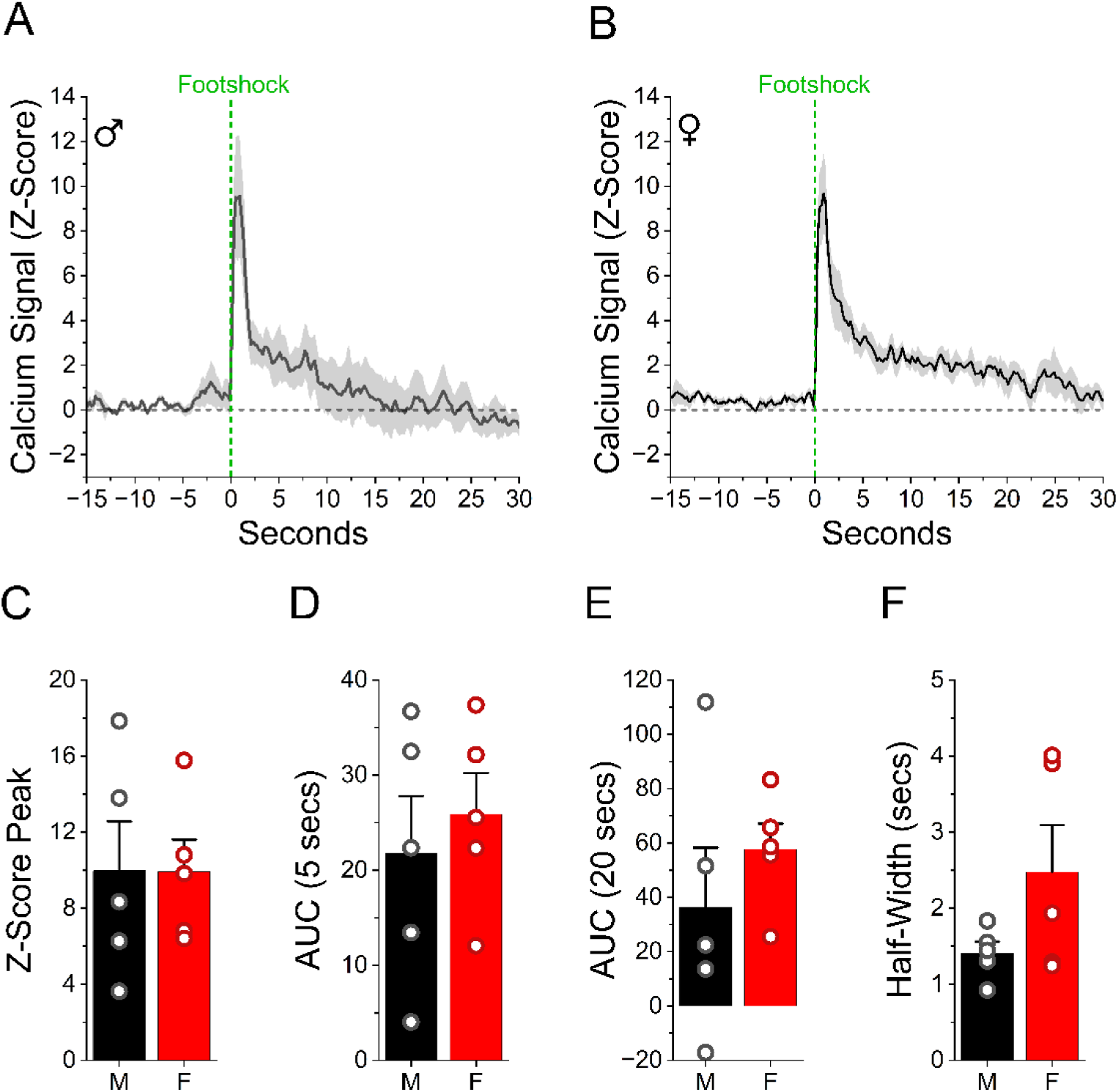
BLA NPY+ Interneurons Respond to Footshock Stimulus in Both Males and Females. A. Calcium fiber photometry trace shows footshock activates NPY cells in male mice (n= 5 hemispheres/4 mice). B. Calcium fiber photometry trace shows footshock activates NPY cells in female mice (n= 5 hemispheres/3 mice). C. Peak Z-score is not different between female and male mice. (Two-tailed Student’s *t*-test, p=0.99). D. AUC (0 to 5 seconds) is not different between female and male mice. (Two-tailed Student’s *t*-test, p=0.60). E. Even a longer AUC (0 to 20 seconds) shows no difference between female and male mice. (Two-tailed Student’s *t*-test, p=0.40). F. The half-width of the calcium signal is not different between male and female mice. (Two-tailed Student’s *t*-test, p=0.13). For all panels the error bars represent ± SEM. * indicates *p* < 0.05.

### 3.5 Differences in BLA NPY+ Interneuron Activation between Mild and Strong Aversive Stimuli

Since air puffs and footshocks both activate BLA NPY+ neurons, this could indicate that these cells are involved in conveying generalized information about any aversive stimulus. However, activation of BLA NPY+ neurons was not to the same extent for both aversive stimuli (n = 10 hemispheres/ 7 mice). Overall, the footshock (FS) protocol showed an enhanced calcium signal of BLA NPY+ cells compared to air puffs (AP). Specifically, the stronger aversive stimulus has a greater effect on the peak Z-score (AP 7.73 ± 1.68 vs FS 9.95 ± 1.45; one-tailed paired *t*-test p=0.037) and AUC (0-20 seconds; AP 23.93 ± 7.88 vs FS 47.12 ± 11.74; one-tailed paired *t*-test p=0.039). Even looking at a shorter window for the AUC, the response is larger with footshock compared to air puff (0-5 seconds; AP 16.70 ± 4.44 vs FS 23.84 ± 3.56; one-tailed paired *t*-test p=0.035). However, the half-width of the calcium signal from BLA NPY+ cells was not significantly different between air puffs and footshocks (AP 1.71 ± 0.30 s vs FS 1.94 ± 0.35 s; one-tailed paired *t*-test p=0.28). Together, these data suggest that the strength of the aversive stimulus modulates the extent of BLA NPY+ cell activation through modulation of either calcium influx and/or recruitment of NPY+ neurons.

### 3.6 No Long-Lasting Changes in BLA NPY+ Neuron Activation in Response to Footshock

Traumatic stress can cause lasting changes to the NPY system; the effects depend on the brain region and type of stressor. Studies using footshock (albeit with different protocols) saw lasting effects on hippocampal but not amygdala NPY expression (Cortes et al., 2021) whereas predator scent stress reduces NPY expression one week after exposure in both regions (Cohen et al. 2012). Predator scent stress abolished NPY release measured in hippocampal slices one week later (Li et al. 2017). Lasting effects of aversive stimuli on *in vivo* NPY+ cell activation have not yet been tested. We next sought to determine whether previous exposure to a strong aversive stimulus would result in changes to BLA NPY+ neuron activation. This required a longitudinal approach employed by re-exposing the same mice to the same footshock protocol 1 week later (Figure 5E). We first determined if this footshock protocol produced any long-lasting behavioral modifications. For this, a separate cohort of mice (without viral injections or fiber placement) were tested for behavioral changes in the open field task 1 week after administration of the footshock protocol (Figure 5A). Mice receiving the footshock protocol traveled shorter distances compared to control mice (Figure 5B), as well as having a reduction in velocity (Figure 5C), indicating a lasting effect of the footshock protocol on locomotion. However, there was no difference in the percentage of distance traveled in the center of the open field (Figure 5D), indicating no overt increase in anxiety-like behavior. Once behavioral changes were verified, we next used a subset of mice from Figure 3 to receive a second session of the footshock protocol (Figure 5E). The GCaMP responses were nearly identical between sessions 1 (S1) and 2 (S2; Figure 5F), indicating no difference in BLA NPY+ neuron activity between the two footshock sessions. There was a slight trend (p=0.07) for the AUC (Figure 5H) to be elevated in response to the second footshock session. But, no changes in Z-Score peak (Figure 5G) or half-width of the calcium signal (S1 2.15 ± 0.48 s vs S2 1.88 ± 0.28 s; one-tailed paired *t*-test p=0.72) were observed in NPY+ cells expressing the GCaMP sensor. Overall, these data demonstrate that a previous physical aversive stimulus does not alter BLA NPY+ neuron activation 1 week later, indicating that NPY+ cell activation is consistent over time.

**Figure 5:**
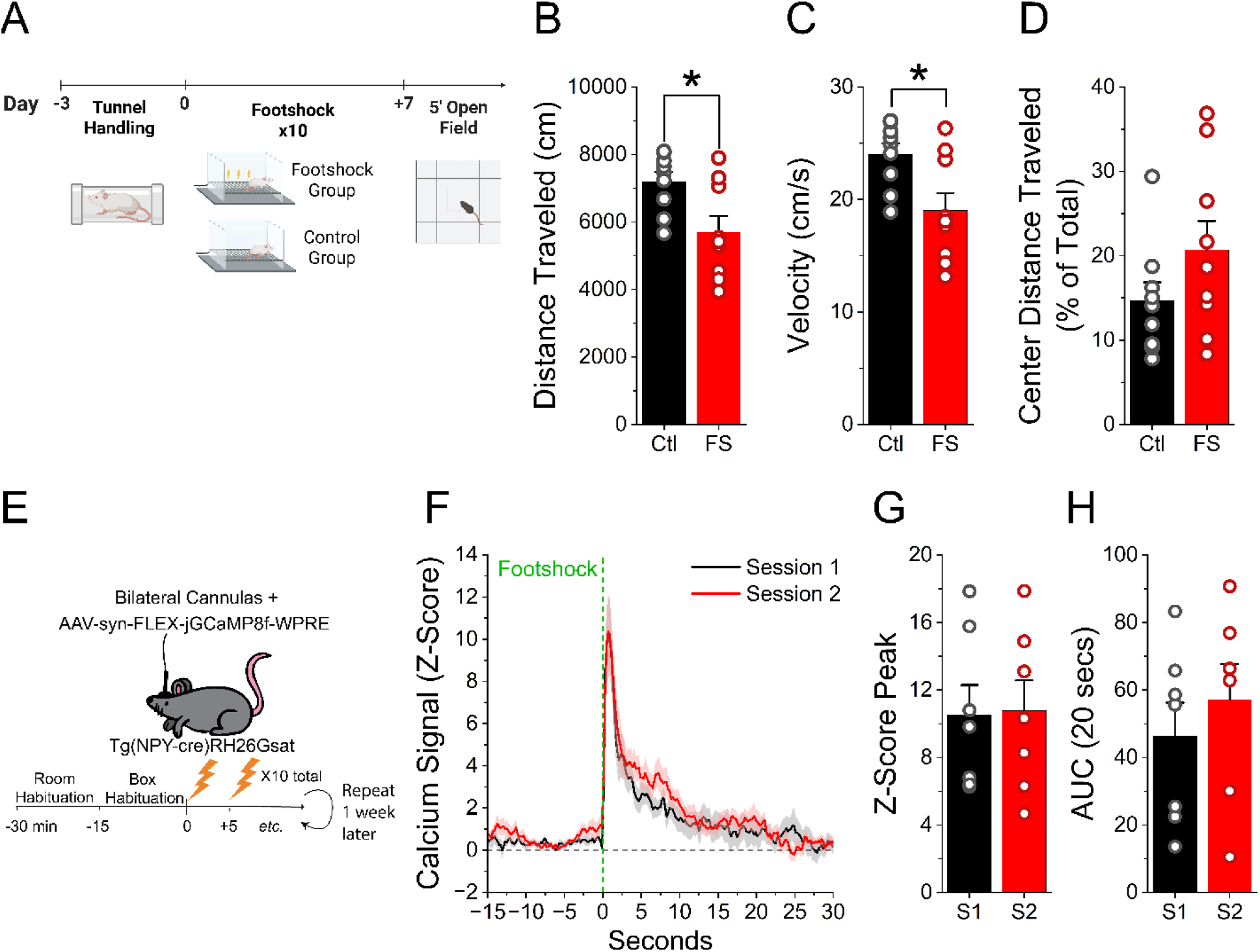
BLA NPY Interneurons Respond Equally to Footshock After 1 Week. A. Schematic of experimental timeline for open field behavioral task. Male and female mice underwent tunnel handling for 3 days. Mice were placed in either a footshock (n=9 mice: 5 F, 4 M) or control (n=9 mice: 6 F, 3 M) group. The footshock group received 10 footshocks (1 sec, 1 mA, 5 minutes apart), whereas the control group was placed in the chamber for the same duration. A week later both groups performed a 5-minute open field behavioral task. B. Total distance traveled is reduced in the footshock (FS) group vs the control (Ctl) group in the open field task. (Two-tailed Student’s *t*-test, p<0.05). C. The footshock group had a slower velocity compared to the control group. (Two-tailed Student’s *t*-test, p<0.05). D. However, the center distance traveled was not significantly different between the two groups. (Two-tailed Student’s *t*-test, p=0.15). E. Surgical design and experimental timeline. Bilateral AAV injection of Cre-dependent GCaMP8f with bilateral insertion of fiber optic cannulas were performed in male and female NPY-Cre^+/−^ mice. Footshock stimulation sessions were administered 1 week apart. Each session consisted of 10 footshocks (1 sec, 1 mA, 5 minutes apart). F. Comparison of calcium response in BLA NPY interneurons between two footshock sessions 1 week apart (n = 7 hemispheres/5 mice: 3F, 2M). G. Peak Z-Score of calcium response is not different between footshock session 1 (S1) and session 2 (S2; One-tailed paired *t*-test, p = 0.32). H. No significant difference was seen in the AUC (0 to 20 seconds) between session 1 and session 2 (One-tailed paired *t*-test, p = 0.07). For all panels the error bars represent ± SEM. * indicates *p* < 0.05.

## 4. DISCUSSION

Here we showed for the first time that aversive stimuli acutely activate amygdala NPY+ cells *in* vivo in both male and female mice. Our data show robust activation of BLA NPY+ neurons during aversive stimuli, indicated by increased GCaMP8f fluorescence in response to air puff and footshock. BLA NPY+ cells are activated to a lesser extent by air puff, a milder stimulus that is uncomfortable but not painful, indicating that NPY+ cells can respond differentially to a variety of aversive stimuli. In both cases, the duration of the calcium signal in NPY+ cells was different between the first and last iteration of the aversive stimuli. However, the duration was diminished with air puff (mild stimulus) and prolonged with footshock (strong stimulus) in response to repeated trials, suggesting subtle differences in the way that NPY+ cells respond to different aversive stimuli. Interestingly, consistent NPY+ cell activation was observed with the footshock paradigm applied one week apart, showing the reliability of the NPYergic system in BLA to aversive stimuli. Our results demonstrate the active involvement of amygdala NPY+ cells in the canonical amygdala-mediated stress circuitry, consistent with the idea that NPY+ cell activation serves as a buffering mechanism, through the release of their neurotransmitters, to regulate stress responses.

The BLA processes aversive stimuli (Beyeler et al., 2016; Correia & Goosens, 2016; Grosso et al., 2018), including discriminating between threatening and safe stimuli (Grosso et al., 2018; Namburi et al., 2016). BLA excitatory pyramidal neurons are known to be highly involved in this process (Kim et al. 2016; Zhang and Li 2018); the role that different subtypes of GABAergic interneurons play during this continues to be uncovered. Our study provides the first direct *in vivo* measurement of BLA NPY+ interneuron responses during stressors, demonstrating that they are activated by both footshock and air puff. Air puff activates around 20% of BLA excitatory neurons in mice (Zhang and Li 2018), while the effects of air puff on BLA inhibitory neurons are not yet known. Footshock modulates activity in the majority of BLA GABAergic cells, with 40% being activated and 40% inhibited (Favila et al., 2025). Another study found that footshock activates parvalbumin-expressing (PV+) and somatostatin-expressing (SST+) cells in BLA (Asim et al., 2024), indicating that multiple GABAergic cell types in BLA process aversive information. In contrast, Wolff et al. found that PV+ and SST+ BLA interneurons are inhibited by footshock during a cued fear conditioning protocol (Wolff et al., 2014), suggesting that the same type of GABAergic interneuron in BLA could respond differently to expected vs unexpected stimuli (Yau et al., 2021). BLA NPY+ cells could respond to both expected and unexpected aversive stimuli, since some activation was observed upon the approach of the air puff canister. Responses from individual BLA NPY+ cells are likely heterogeneous, as observed for other BLA inhibitory neuron subtypes (Favila et al., 2025), yet the overall effect leads to enhanced activation of NPY+ cells. Interestingly, BLA NPY+ interneurons do not respond to all aversive stimuli, as c-Fos levels were unchanged following elevated plus maze exposure (Butler et al., 2012; Regev-Tsur et al., 2020). Our results shed new light on another GABAergic cell subtype engaged during the presentation of an aversive stimulus in the BLA.

The neural encoding of aversive events involves rapid activation of threat detection systems, primarily centered in the amygdala, which then orchestrate widespread changes in brain activity and peripheral physiology (LeDoux, 2003; Mahan & Ressler, 2012; Steinberg et al., 2020). Multiple brain regions showed that GABAergic interneurons respond to aversive stimuli (Li et al. 2024; Allen et al. 2017), and this holds true for NPY+ cells. Stressors have been shown to activate NPY+ interneurons in the dorsal raphe nucleus (Zhang et al. 2024), a hub for serotonin-expressing neurons, and locus coeruleus (Riga et al., 2025), a primary source of norepinephrine. Activation of NPY+ cells in these brain regions could modulate the serotonin and/or norepinephrine response to a stressor. Similarly, NPY+ neurons could be involved in modulating coping strategies during a stressor, as NPY+ cells responded to an aversive stimulus in the ventrolateral periaqueductal grey regions (Zhang et al. 2024), a brain region important for passive coping strategies. BLA principal neurons encode valence-specific information, with distinct populations responding to aversive versus appetitive stimuli (Beyeler et al., 2016). This valence coding depends critically on the balance between excitation and inhibition provided by local interneuron populations, and the NPY+ neurons in BLA that we studied could play a role in assigning the valence. Additionally, BLA NPY+ cells could convey generalized information about aversive events, since both stressors activated the cells. However, there were subtle differences in the extent of the response between the two stimuli, suggesting that NPY+ cells could still be involved in the discrimination of different aversive stimuli. Importantly, our results demonstrate that BLA NPY+ cells respond to aversive stimuli in the central hub of the threat detection system.

Our study provides the first real-time temporal characterization of *in vivo* BLA NPY+ interneuron activity. The majority of previous studies have used c-Fos to identify activation of NPY cells in response to aversive stimuli (Regev-Tsur et al., 2020; Riga et al., 2025); its fixed time window limits the understanding of cell activation dynamics. Using in vivo calcium imaging, we show that NPY+ cell activation occurs rapidly, as the peak responses occurred within milliseconds, and repeatedly, as responses were seen with each iteration of either aversive stimulus. While both footshock and air puff activated BLA NPY+ cells, the stronger aversive stimuli caused a larger calcium response compared to the weaker stimuli. This could be due to a larger calcium influx per cell, and/or recruitment of greater numbers of BLA NPY+ cells in response to footshock. Another possibility is that different subpopulations of BLA NPY+ cells may respond to specific stimuli, as has been seen for other BLA neurons (Grosso et al., 2018; Liu et al., 2021). Odorant air puffs activate parvalbumin-expressing and somatostatin-expressing, however, the kinetics differ between the two cell types (Allen et al., 2017). Somatostatin-expressing cells had a slower onset and decay compared to parvalbumin-expressing cells (Allen et al., 2017). We find that BLA+ NPY cells have a fast onset to both aversive stimuli (< 1 second), yet there was a broad range in latency of the peak maximum. Subpopulations of NPY+ cells also express somatostatin (McDonald, 1989; Sosulina et al., 2006), potentially explaining the variability in onset kinetics. Interestingly, the response duration to repeated trials was longer with the stronger stimulus (footshock), suggesting that this stimulus elicits more action potential firing. Activation of NPY+ interneurons would be expected to initially release GABA, and with higher frequency firing, there could be release of NPY (Li et al. 2017) as well. It is unknown if the level of activation observed in this study would induce NPY release for either aversive stimulus, but NPY release to footshock was seen in the locus coeruleus (Riga et al., 2025). Because both GABA and NPY cause dampening of circuits, this could result in both rapid (through GABA release) and sustained (through NPY release) anxiolytic effects in response to traumatic stressors. Being able to monitor activation of the NPYergic system in real time can provide critical insights into the threat-response dynamics of NPY+ cell activity.

The role of specific GABAergic subtypes has been more widely studied using chronic stress paradigms (Albrecht et al., 2021). Previous studies using c-Fos showed that NPY+ cells are activated by prolonged stress (multiple days) in the hippocampus (Regev-Tsur et al., 2020) and acute novelty stress (2 hours) in the dorsal raphe nucleus and ventrolateral periaqueductal gray region (Zhang et al. 2024). Our data show that prolonged stress exposure is not required, as BLA NPY+ cells are acutely activated by a single application of either air puff or footshock. Other GABAergic interneurons in BLA have shown reduced activity (habituation) in response to repeated footshocks (Asim et al., 2024). Additionally, habituation activity to air puff has been observed in heterogeneous populations of glutamatergic and GABAergic cells in other brain regions (van der Zouwen et al., 2025). In contrast, BLA NPY+ cells continue to be activated by multiple presentations of either stimulus, with no change in the peak amplitude. However, we did see changes in the duration of response of BLA NPY+ cells which differed between footshock and air puff. In particular, footshock caused an increased duration of NPY+ cell responses in later iterations, while air puff caused reduced duration. Our data show for the first time that BLA NPY+ cells respond to acute stressors with varying patterns of activity during repeated iterations.

Interestingly, two footshock sessions a week apart produced consistent BLA NPY+ interneuron activation, highlighting the resilience of the NPYergic system in the amygdala. Despite eliciting aversive-event responses, the protocol did not cause lasting changes in NPY+ cell activation, indicating the stressor was below the threshold for maladaptation seen in other stress protocols (Cohen et al. 2012; Li et al. 2017; Cortes et al. 2021). Reduced NPY expression in amygdala was observed with predator scent stress (Tzanoulinou et al. 2014; Cohen et al. 2012) but not with unpredictable footshock (Cortes et al., 2021), although both reduced hippocampal NPY expression (Cohen et al. 2012; Cortes et al. 2021). The footshock protocol used here could therefore be used to probe NPY+ amygdala circuitry dynamics in response to other interventions, such as PTSD protocols. Higher-intensity or chronic stress paradigms might reveal lasting changes in NPY+ cell activation, as NPY modulates stress resilience (Reichmann & Holzer, 2016; Silveira Villarroel et al., 2018). Other studies revealed that stress protocols altered GABA system function (Tzanoulinou et al., 2014) and/or changes in GABA synaptic responses (Isoardi et al., 2007), suggesting long-lasting effects on interneuron activation not observed here. The stable response of NPY+ cells to repeated stressors supports NPY’s proposed role as a buffering mechanism (Michaelson et al., 2020), potentially preventing overexcitation of amygdala circuits under acute stress.

The NPY+ cell activation peak response to both stimuli had a large range in both male and female mice. All included hemispheres showed expression in BLA; the majority showed only BLA expression, but a minority showed a mix of BLA and central amygdala (CeA). As a result, the distribution of GCaMP expression between CeA and BLA could potentially play a role in the variability of the response. Hemispheres only with expression in BLA had a peak response to footshock that ranged from 3.6 to 17.8, whereas hemispheres with expression in both BLA and CeA ranged from 6.4 to 15.8. Therefore, it is unlikely that the variability is due to differences between BLA and CeA responses. Instead, the variability in peak responses is probably due to differences in GCaMP expression levels and/or the distance of the sensor from the fiber tip.

Here we demonstrate the first real-time measurement of BLA NPY+ neuron activation *in vivo*, illustrating that these cells respond to aversive stimulation such as footshock. Determining when and where NPY+ cells are activated, and thus potentially triggering NPY release, is crucial for understanding NPY’s role in normal brain function and disease. Understanding disease-related changes to NPY+ cell activity could open new avenues for exploring NPY’s therapeutic potential in stress-related disorders.

## Notes

### Competing Interest Statement

The authors have declared no competing interest.

### Funding

- National Institute of Mental Health: R01MH108342

